# Analysis of crosslinking sites suggests *C. elegnas* PIWI Argonaute exhibits flexible conformations for target recognition

**DOI:** 10.1101/2025.02.14.638322

**Authors:** Wei-Sheng Wu, Dong-En Lee, Chi-Jung Chung, Shang-Yi Lu, Jordan S. Brown, Donglei Zhang, Heng-Chi Lee

**Affiliations:** Department of Electrical Engineering, National Cheng Kung University, Tainan 701, Taiwan; Department of Molecular Genetics and Cell Biology, University of Chicago, Chicago, IL 60637, USA

## Abstract

Small RNAs play critical roles in gene regulation in diverse processes across organisms. Crosslinking, ligation, and analyses of sequence hybrid (CLASH) experiments have shown PIWI and Argonaute proteins bind to diverse mRNA targets, raising questions about their functional relevance and the degree of flexibility in target recognition.

As crosslinking-induced mutations (CIMs) provides nucleotide-resolution of RNA binding sites, we developed MUTACLASH to systematically analyze CIMs in piRNA and miRNA CLASH data in *C. elegans*. We found CIMs are enriched at the nucleotide positions of mRNA corresponding to the center of targeting piRNAs and miRNAs. Notably, CIMs are also enriched at nucleotides with local pairing mismatches to piRNA. In addition, distinct patterns of CIMs are observed between canonical and non-canonical base pairing interactions, suggesting that the worm PIWI Argonaute PRG-1 adopts distinct conformations for canonical vs. non-canonical interactions. Critically, non-canonical miRNA or piRNA binding sites with CIMs exhibit more regulatory effects than those without CIMs, demonstrating CIM analysis as a valuable approach in assessing functional significance of small RNA targeting sites in CLASH data. Together, our analyses reveal the landscapes of Argonaute crosslinking sites on mRNAs and highlight MUTACLASH as an advanced tool in analyzing CLASH data.

## Introduction

Distinct types of small RNAs, such as miRNAs and piRNAs, are critical regulators of gene expression in diverse animals^1^. Small RNAs guide Argonaute family proteins to recognize mRNA targets through base pairing interactions between small RNAs and mRNAs^12^. Specifically, pairing between the seed region, the second to the seventh or eighth nucleotide of small RNAs, and the target mRNA is reported to play a critical role in target recognition of animal miRNA and piRNAs^34^. However, both canonical (perfect seed pairing) and non-canonical (imperfect seed pairing) base pairing interactions can result in gene silencing, prediction of functional small RNA targeting sites is difficult^5,6^. Crosslinking, ligation, and analysis of sequence hybrid (CLASH) is a critical experimental approach for the identification of small RNA binding sites *in vivo*^7^. In CLASH experiments, UV crosslinking leads to the formation of covalent bonds between Argonautes and their interacting mRNAs, allowing for a stringent purification process. The small RNAs are then ligated to partially degraded target mRNAs, which are then cloned into cDNA libraries and sequenced. Analysis of hybrid reads consisting of a small RNA and its target RNA can, therefore, reveal the transcriptome-wide interactions between small RNAs and their target mRNAs^7^. Surprisingly, the majority of hybrid reads from CLASH data do not contain canonical small RNA-mRNA interaction^7–9^. Further analyses of miRNA binding sites with non-canonical binding suggest that most of these non-canonical binding sites events exhibit little functional relevance^10^. At the same time, empirical studies of small RNA-mediated gene regulation have demonstrated that non-canonical small RNA binding sites can be of functional significance^5,11,12^. Currently, it is difficult to evaluate which of the non-canonical binding sites identified in CLASH data are of functional significance.

Crosslinking-induced mutations (CIMs) are produced upon cDNA synthesis during CLIP (crosslinking immunoprecipitation) experiments, where CIMs reveal the footprints of distinct RNA binding proteins on their target mRNAs^13^. While there are several tools to analyze CLASH data^14–17^, they do not consider CIMs. Here, we developed the MUTACLASH analysis pipeline to identify CIMs data and examined CIMs from piRNA and miRNA CLASH data in *C. elegans*. Our results suggest that CIMs are enriched at the center of the small RNA binding sites. In addition, CIMs from piRNA targeting sites are also enriched at nucleotides with local mismatches. In particular, we observed that target sites with canonical or non-canonical piRNA-mRNA basepairing are associated with distinct crosslinking patterns, suggesting that worm PIWI Argonaute PRG-1 adopts distinct conformations for canonical and non-canonical target recognition. Importantly, piRNA and miRNA binding sites with CIMs generally exhibit stronger regulatory effects than those without CIMs, including those without canonical base pairing interactions. Together, our CIMs analyses of CLASH data reveal the crosslinking landscape of piRNA PIWI and miRNA Argonaute on their target mRNAs and provide insight into the target recognition by PIWI family proteins. Our results using MUTACLASH further demonstrate that CIMs analysis offers critical information in evaluating the functional significance of CLASH-identified small RNA binding sites.

## Results

### Identification of CIMs from PIWI and Argonaute CLASH data

As mentioned above, CIMs represent the footprints of RNA binding proteins on their target mRNA^13^. Since both CLIP and CLASH experiments involve UV crosslinking, we reasoned that CIMs should also be present in PIWI/Argonaute CLASH data (Figure 1A) and those crosslinking sites likely represent the footprint of PIWI/Argonaute on their target mRNAs as well.

**Figure 1.**
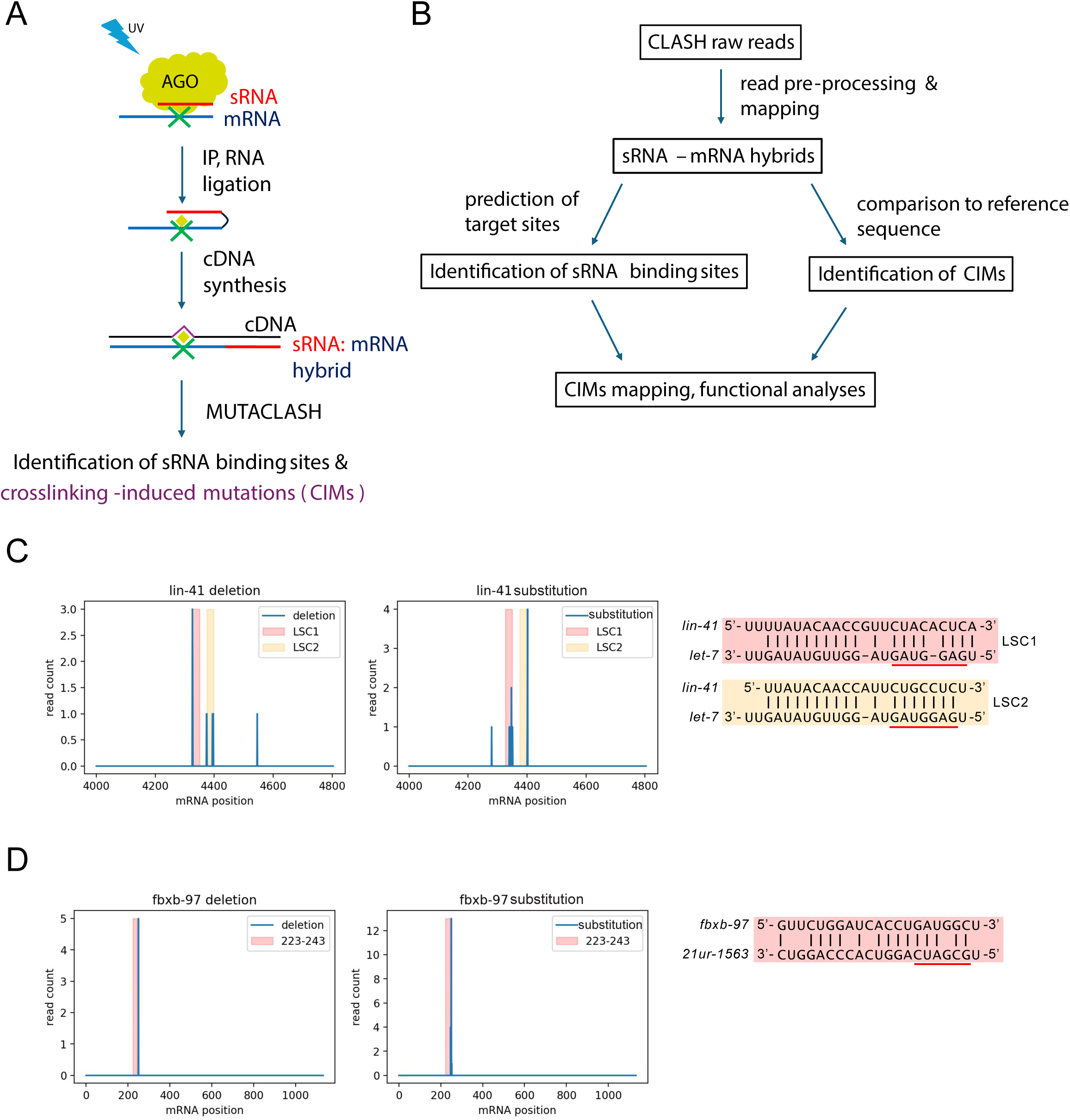
Identification of Crosslinking-Induced Mutations (CIMs) in CLASH Data at piRNA and miRNA Targeting Sites. A. A model illustrating the CLASH experimental procedure of PIWI and Argonaute complexes and the production of CIMs during cDNA synthesis. B. A flowchart of MUTACLASH analysis pipeline for identifying small RNA-mRNA hybrids and for analyzing CIMs from PIWI and Argonaute CLASH data. C. The distribution of deletions (left) and substitutions (middle) on *lin-41* mRNA identified from hybrid reads with the *let-7* miRNA from ALG-1 Argonaute CLASH data. The base-pairing between *let-7* miRNA binding sites, LSC1 and LSC2, on the *lin-41* 3’ UTR is shown (right). The seed region of the let-7 miRNAs is underlined. D. The distributions of deletions (left) and substitutions (middle) on *fbxb-97* mRNAs identified from hybrid with piRNA *21ur-1563* from PRG-1 PIWI CLASH data. The base-pairing between piRNA *21ur-1563* and mRNA *fbxb-97* is shown (right). The seed region of the piRNA is underlined.

To examine whether the CIMs, such as mRNA deletion or substitution, are present in CLASH data, we developed the MUTACLASH analysis pipeline that take raw sequencing reads from CLASH data that integrate those previous published algorithms to identify small RNA-mRNA hybrids^14–17^ and critically, to locate CIMs within these hybrids (Figure 1B). We first examined hybrids derived from previously well-characterized piRNA and miRNA targeting sites, including *lin-41* mRNA (targeted by miRNA *let-7*) and *fbxb-97* mRNA (targeted by piRNA *21ur-1563*)^8,9,11^. In those target mRNAs, we found that CIMs, including both deletions and substitutions, are present in hybrid reads of those targeting sites (Figure 1C and 1D). Notably, CIMs are present at both the canonical target sites with perfect seed pairing, such as the *let-7* miRNA target site LSC2 on *lin-41* miRNA, as well as the non-canonical target sites with seed mismatch or bulge, such as the *let-7* miRNA target site LSC1 on *lin-41* mRNA, or the *21ur-1563* piRNA target site on *fbxb-97* mRNA (Figure 1C and 1D). These observations supported that CIMs are present at piRNA PIWI and miRNA Argonaute CLASH data.

We then applied MUTACLASH to systematically analyze the presence of CIMs in the PIWI PRG-1 (piRNA) CLASH and Argonaute ALG-1 (miRNA) CLASH/iCLIP data^8,9^. To assess whether these mutations identified from MUTACLASH are indeed induced by crosslinking, we first compared the mutations of mRNAs from CLASH reads with that from mRNA sequencing reads, which are obtained with and without UV crosslinking in their experimental procedures, respectively. Our analysis indicated that mRNA derived from PRG-1 and ALG-1 CLASH data exhibited a higher incidence of deletions (6.55% and 5.8%) or substitutions (6.67% and 7.41%) compared to deletions (0.026%) and substitutions (1.4%) from mRNA sequencing data (Table S1A). While the percentage of mRNA segments in hybrid reads containing CIMs is lower than that in mRNA (non-hybrid) reads from PRG-1 CLASH data, it remains significantly higher than what is observed in mRNA sequencing data (Table S1A).

In addition, we observed that CIMs identified from PRG-1 or ALG-1 CLASH data, both non-hybrid and hybrid reads, exhibited a preference for uridine-derived mutations (ranging from 41% to 61%) (Table S1B). On the contrary, no such uridine preference was found in mRNA sequencing data. These findings align with previous reports that uridine is preferentially crosslinked to proteins during UV crosslinking^18,19^. Moreover, we observed a strong preference for specific types of substitutions in PRG-1 and in ALG-1 CLASH data, including U to C substitutions (representing 79-90% of all U substitutions) and C to U substitutions (representing 68-84% of all C substitutions) (Table S1C and S1D). Collectively, the elevated mutation frequency and mutation preference confirmed that CIMs are widely present in CLASH data.

### CIMs are enriched at nucleotides corresponding to the center of piRNA and miRNA binding sites

Previous analyses of CIMs for CLIP experiments of various RNA-binding proteins have shown that CIMs are frequently located at specific positions within their mRNA binding motifs^20^. Unlike sequence-specific RNA-binding proteins, PIWI and Argonaute family proteins are guided by distinct small RNAs to target mRNAs. CLIP analysis of neuronal tissue has implied that CIMs may occur in the middle of the miRNA pairing site^13^. However, since CLIP data only reveal the identity of mRNAs but lack information about bound miRNAs, the pairing information and the location of CIMs can only be inferred from CLIP analyses. Therefore, it remains unclear whether CIMs are enriched at specific positions of PIWI and Argonaute binding sites. CLASH data offer the ideal dataset to examine the distribution of CIMs in Argonaute binding sites as hybrid reads provide both the identity of mRNAs and targeting miRNAs^7^. We applied MUTACLASH to analyze PRG-1 piRNA CLASH data and to calculate the distribution of CIMs at single nucleotide resolution across the entire piRNA targeting sites^8^. We found that CIMs detected in mRNAs were located preferentially at nucleotides corresponding to the center of piRNA-mRNA interactions (Figure 2A). Specifically, mRNA deletions were peaked at positions corresponding to the 11th and 12th nucleotides of piRNAs, while the mRNA substitutions were peaked at positions corresponding to the 9th and 10th nucleotides of piRNAs (Figure 2A). This enrichment of CIMs at the center region was even more pronounced when we only consider those piRNA target sites within the top 33% piRNA targeting scores^21^ (Figure S1A). In these sites with high piRNA targeting scores, a minor peak of CIMs can also be found at the beginning of piRNAs.

**Figure 2.**
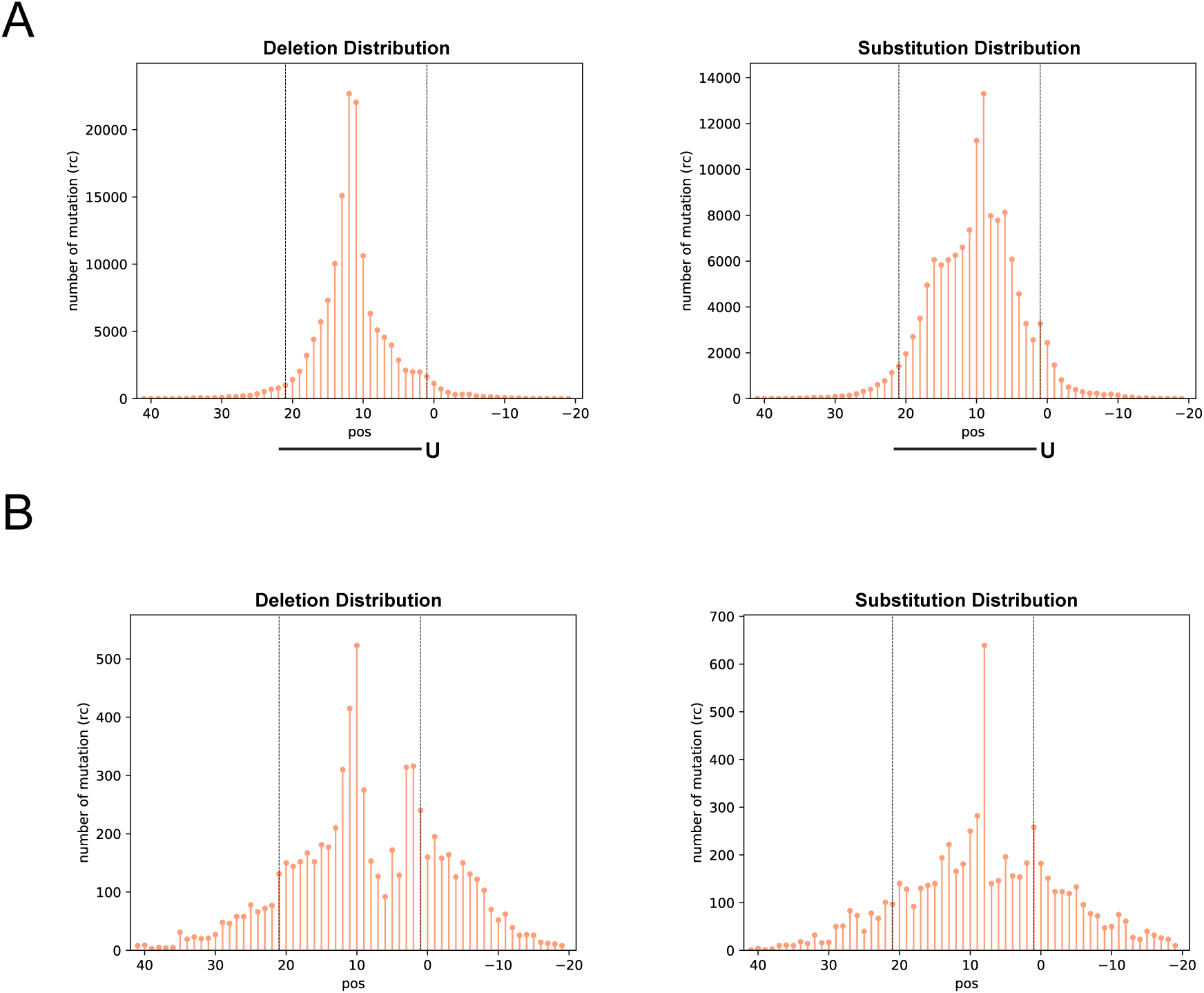
The distribution of CIMs of at piRNA and miRNA targeting sites. A. The number of deletions (left) and substitutions (right) found at the indicated position of mRNAs corresponding to their targeting piRNAs from PRG-1 piRNA CLASH data. B. The number of deletions (left) and substitutions (right) found at the indicated position of mRNAs corresponding to their targeting miRNA ALG-1 miRNA Argonaute CLASH data.

We then analyzed the position of CIMs in miRNA (ALG-1 Argonaute) CLASH data of *C. elegans*^9^. Interestingly, several of the observed in in piRNA CIM distributions were also be found in miRNAs; First, CIMs were also enriched at nucleotides corresponding to the center and slightly at the beginning of their target sites (Figure 2B). The peak positions of deletions and substitutions are also enriched at 11/12^th^ and 9/10^th^ position of miRNA, respectively.

Furthermore, when considering only sites within the top 33% of miRNA targeting scores (MiRanda >130)^22^, a more pronounced peak of CIMs can also be found at position corresponding to the beginning of miRNAs (Figure S1B).

As described above, there is a ∼2 nucleotide shift between the peak position of crosslinking-induced deletions and substitutions in piRNA and miRNA CLASH data. To further examine whether this trend can be observed from the identical sets of small RNA targets, we limited our analysis to those hybrids where both substitutions and deletion were found at the same small RNA targeting sites. We found that position of deletion preferentially occurs mostly at two nucleotides upstream of the position of substitution on mRNAs (Table S2A).

Together, our analyses suggest that the CIMs are preferentially produced on piRNA or miRNA targets at regions corresponding to the center of the small RNAs, and in some cases also at the beginning of the small RNAs. The consistency in CIM distribution between these unique small RNA targeting events supports the notion that these CIMs act as the footprints of PIWI/Argonaute on their target mRNAs^20^.

### CIMs are also enriched at regions of piRNA target sites with local mismatches

Base-pairing at the seed region, which is the 2^nd^ to 7^th^ or 8^th^ nucleotides of piRNA or miRNA, is known to play a critical role in piRNA and miRNA targeting in *C. elegans*^4,9^. As we noticed that CIMs are somewhat depleted in their seed regions (Figure 2A and 2B), we wondered whether the local base-pairing between small RNAs with mRNAs could affect the distribution of CIMs. To examine the relationship between CIM position and base-pairing, we combined all hybrids with CIMs corresponding to each piRNA position from PIWI CLASH data and calculated the base-pairing ratio for each position, then compared those ratios. Notably, we noticed a trend that the piRNA base-pairing ratio is reduced around the location of the CIMs; For example, hybrids with mRNA deletion or substitution at the position corresponding to the 5^th^ nucleotide of the piRNA exhibits a reduced pairing ratio around the 5^th^ nucleotide when compared to all hybrids (Figure 3A). Similarly, hybrids with CIMs at the 14^th^ nucleotide exhibit a reduced pairing ratio around 14^th^ position (Figure 3B). In addition, we noticed that the overall base-pairing ratio at seed and non-seed regions seems associated with whether the CIMs are located in seed or non-seed region (see more detailed analyses in the next session). This global trend of reduced base-pairing ratios near the location of CIMs can be further demonstrated when we aligned all CIMs at the center (position 0) and compared the pairing ratio around the CIMs. Indeed, we found that the base-pairing ratios were reduced around the location of CIMs (Figure 3C). However, in ALG-1 miRNA CLASH data, such trends could only be clearly found in the substitution, but not the deletion, carrying hybrids (Figure S2A). Our analyses reveal that CIMs preferentially occur at nucleotides of piRNA targeting sites with less stable local base-pairing.

**Figure 3.**
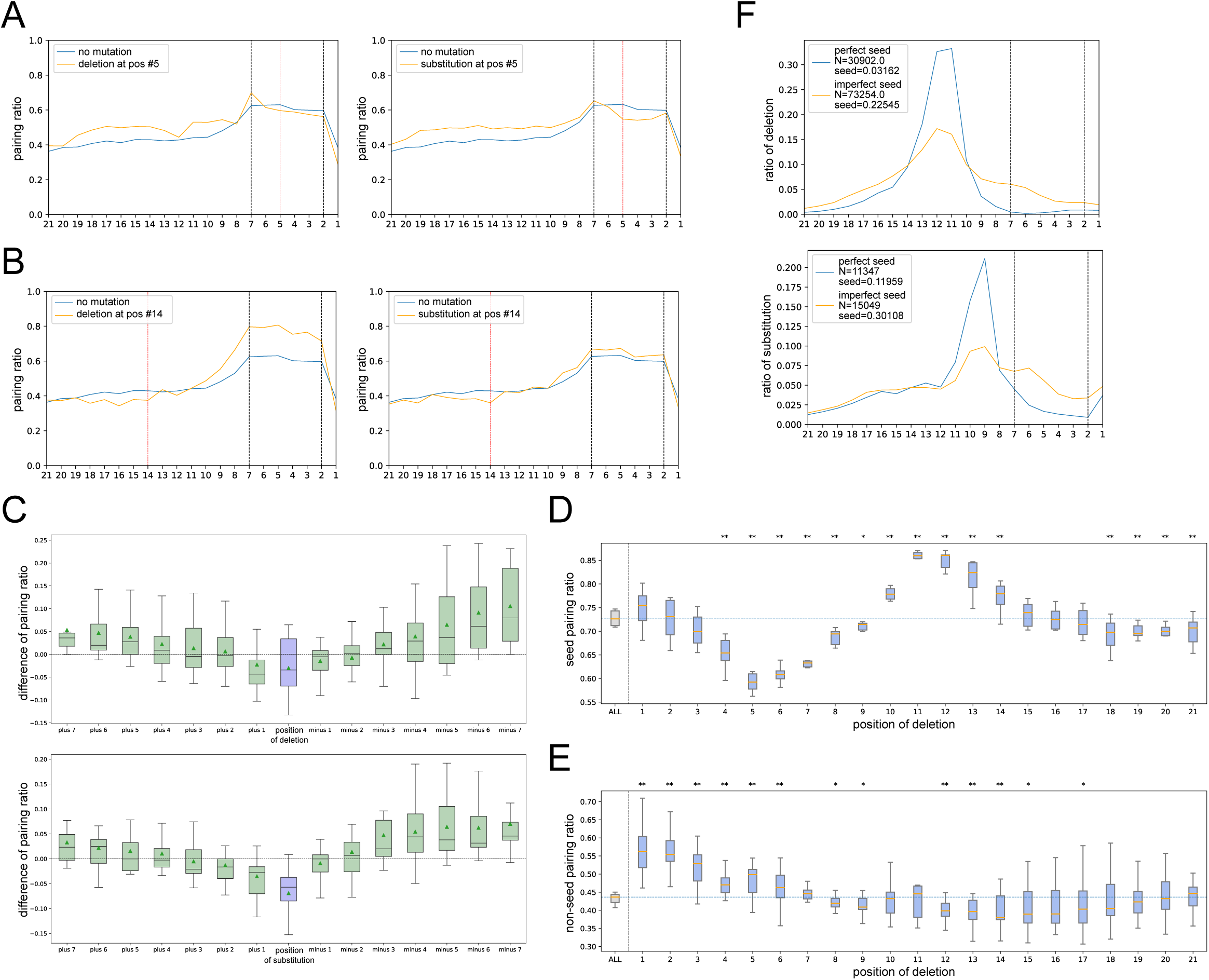
CIMs preferentially occur at positions with local piRNA pairing mismatches. A. Pairing ratios between mRNA and piRNA at the indicated positions from hybrids containing mRNA deletions (left) or substitutions (right) at position 5 (red line) compared to hybrids without mutations (orange lines) in PRG-1 PIWI CLASH data. B. Pairing ratio of mRNA and piRNAs at the indicated position of piRNAs from hybrids containing mRNA deletion (left) or substitution (right) at the position 14 (red line) compared to hybrids without mutations (orange lines) in PRG-1 PIWI CLASH data. C. Differences in base-pairing ratios around CIMs in PRG-1 PIWI CLASH data. The differences were calculated by subtracting the pairing ratios of hybrids with deletions (top) or substitutions (bottom) from the pairing ratios of all hybrids. The analysis was centered on the CIM location (position 0) and extended to the indicated upstream and downstream nucleotides. D. Differences of pairing ratio at the seed region between hybrids with deletion at the indicated position and hybrids with CIMs in PRG-1 PIWI CLASH data. E. Differences of pairing ratio at the non-seed region between hybrids with deletion at the indicated position and hybrids with CIMs in PRG-1 PIWI CLASH data. F. Distribution of mRNA deletions (left) and substitutions (right) in hybrids with either perfect seed pairing (blue line) or imperfect seed pairing (orange line) in PRG-1 PIWI CLASH data.

### Canonical and non-canonical piRNA target sites exhibit distinct CIM pattern

As mentioned above, we noticed a relationship between CIM locations and the overall piRNA paring ratio at seed and non-seed regions: when CIMs occur at non-seed regions, such as position 14, we observed an overall increased pairing ratio at nucleotidess within the seed region, accompanied by slightly decreased or no-change paring ratios at nucleotide of the non-seed region (Figure 3B). On the contrary, when CIMs occur at the seed region, such as position 5, these target sites exhibit an increased paring ratio at nucleotides in the non-seed region, but a decreased or no change in pairing ratios at nucleotides within the seed region (Figure 3A). We then systematically compared the location of CIM position and overall piRNA pairing ratio at the seed region (position 2-7) or non-seed region (8-21). We found that hybrids with CIMs at some of the non-seed positions, such as those with mRNA deletion at the position corresponding to the 11^th^ and 12^th^ of piRNAs or mRNA substitution at the position corresponding to the 1^st^, 9th and 10^th^ of piRNA, exhibit the highest pairing ratios at the seed region when compared to all hybrids carrying CIMs (Figure 3D, Figure S2B). On the contrary, we noticed that hybrids with mRNA deletion or substitution at several positions within the seed positions exhibit a reduced pairing ratios at the seed region but also an increased pairing ratio at the non-seed region (Figure 3D, 3E and S2B, S2C). These observations suggested that CLASH hybrids with CIMs at the center region are associated with sites with a higher seed pairing ratio and thus may be enriched for canonical piRNA target sites. On the contrary, hybrids with CIM at the seed regions are associated with sites with reduced seed pairing ratio and thus may enrich for non-canonical piRNA target targets. Notably, in these non-canonical sites, the reduced seed pairing ratios were accompanied by elevated base-pairing ratios at the non-seed regions (Figure 3E and Figure S2C), likely necessary to establish piRNA binding.

For ALG-1 miRNA CLASH data in *C. elegans*, we also observed a decreased pairing ratio at the seed regions for those hybrids with CIMs located at several positions within the seed region position, such as positions between 4^th^ and 7^th^ (Figure S2D, S2E), but we did not observe a clear increase of non-seed pairing for those hybrids (Figure S2D, S2E). Somewhat similar to piRNAs, hybrids with deletions and substitutions in the middle region exhibited a slightly increased base-pairing ratio at the seed region (Figure S2D).

These observations of the relationship between CIM position and the base-pairing ratios in the seed and non-seed regions of piRNA CLASH data suggest that the PRG-1 PIWI Argonaute protein may adopt distinct conformations in recognizing seed perfect or seed imperfect targets, resulting in distinct crosslinking patterns. If so, hybrids with perfect and imperfect seed base-pairing should exhibit distinct CIM distributions. Consistent with our hypothesis, we found that target sites with perfect seed base-pairing showed stronger enrichment of CIMs at the central region, whereas sites with imperfect seed base-pairing exhibited a more dispersed CIMs distribution, with a greater proportion located in the seed region (Figure 3F). For miRNA ALG-1 targeting sites, those with imperfect seed pairing exhibit more CIMs in the seed region, despite the changes being much less pronounced in sites with mRNA substitutions (Figure S2F).

Together, we found that PIWI Argonaute PRG-1targets exhibit distinct CIMs pattern between perfect seed or imperfectly seed pairing. These observations support the model that PIWI PRG-1 adopt distinct structural conformations for canonical (perfect seed) vs. non-canonical (imperfect seed) targeting, leading to distinct patterns of CIMs.

### Target sites with CIMs exhibit stronger regulatory effects

Previous analyses have reported that the majority of hybrids identified from piRNA and miRNA CLASH data exhibit various base-pairing modes^8,9,23^. Nonetheless, the available piRNA or miRNA targeting score algorithms mainly reward canonical seed perfect base-pairing^21,22^.

When we analyzed the *C. elegans* piRNA and miRNA targets identified through CLASH data (Figure 4A), we found that most hybrids do not contain canonical piRNA or miRNA regulatory sites (pirScan score >0, or miRanda score > 140).

**Figure 4.**
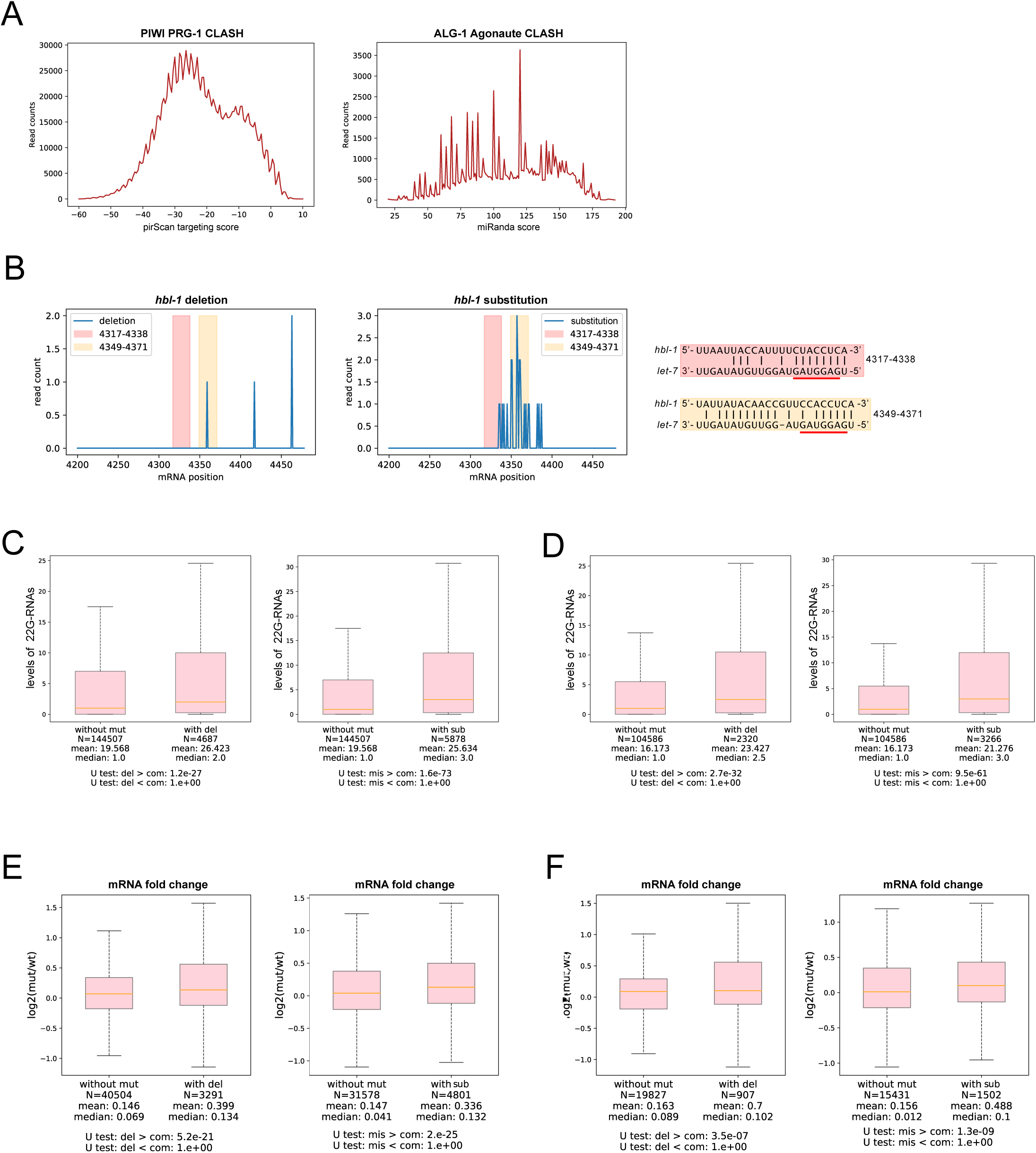
small RNA targets with CIMs exhibit stronger regulatory effects. A. Distribution of piRNA targeting score (pirScan) and miRNA targeting score (miRanda) from the hybris of PRG-1 piRNA CLASH data (left) and ALG-1 miRNA CLASH data (right), respectively. B. Distribution of deletions (left) and substitutions (right) on *hbl-1* mRNA identified from hybrid reads with the *let-7* miRNA from ALG-1 Argonaute CLASH data. The base-pairing between *let-7* miRNA binding sites and two of its targeting sites on *hbl-1* 3’ UTR is shown (bottom). C. Comparison of local WAGO-1 22G-RNAs levels of piRNA targeting sites between those without mutation and those with deletion (left) or substitutions (right). The 22G-RNA levels mapped within the 100-nucleotide window centered at the piRNA targeting site were calculated. The statistical significance (P-value) of the difference in fold change was calculated by the Mann-Whitney U test. D. Comparison of local WAGO-1 22G-RNAs levels of non-canonical piRNA targeting sites between those without mutation and those with deletion (left) or substitution (right). The 22G-RNA levels that mapped within the 100 nucleotide window centered at the piRNA targeting site were calculated. The statistical significance (P-value) of the difference in fold change was calculated by the Mann-Whitney U test. E. The ratio of mRNA level in the *alg-1* mutant over those in wild type of all miRNA targeting sites without CIMs and of those sites with deletion (left) or substitution (right). The statistical significance (P-value) of the difference in fold change was calculated by the Mann-Whitney U test. F. The ratio of mRNA level in the *alg-1* mutant over those in wild type of non-canonical miRNA targeting sites (miRanda score<100) without CIMs and of those sites with deletion (left) or substitution (right). The statistical significance (P-value) of the difference in fold change was calculated by the Mann-Whitney U test.

It was reported that most sites with non-canonical binding sites have little regulatory effects^10^. At the same time, some non-canonical interactions of miRNA and piRNA targeting sites have been shown to elicit gene regulation^5,11,12^, highlighting the importance of developing approaches to reveal non-canonical targets with potential functional significance. Since hybrids from several functional non-canonical miRNA/piRNA sites contain CIMs (Figure 1C), we hypothesized that the CLASH identified small RNA target sites with CIMs are enriched for functional sites. To test this, we first examined *hbl-1*, a known mRNA target of *let-7* miRNA with several predicted *let-7* binding sites in its 3’ UTR^24^. We identified CIMs around *let-7* target sites on *hbl-1* mRNA. Interestingly, while comparing two adjacent *let-7* targeting sites on *hbl-1*, we found more deletions and substitutions at the site containing a seed mismatch than the adjacent site containing a perfect seed match (Figure 4B). Notably, while the contribution of these sites on *hbl-1* gene regulation has not been directly tested, the non-canonical, seed imperfect site has been reported to be the best evolutionally conserved site among all *let-7* target sites on *hbl-1*^24^, implying its functional significance.

Encouraged by these observations, we reasoned that hybrids with CIMs may be enriched for piRNA targeting sites with functional significance. We compared the transcriptome-wide regulatory effects of hybrids with or without CIMs. In *C. elegans*, piRNAs induce gene silencing of their target through the production of secondary small RNAs, known as WAGO 22G-RNAs, locally at piRNA targeted sites^25,26^. We therefore used the local WAGO 22G-RNA levels at piRNA target sites as a proxy to evaluate the functional relevance of piRNA targeting sites. In this analysis, we limit our analysis of WAGO 22G-RNAs mapped to germline-silenced mRNAs (WAGO targets), which are known to be regulated by piRNAs^4,27,28^. We found there are significantly more WAGO-1 22G-RNAs produced around piRNA targeting sites from those hybrids with CIMs than from those without (Figure 4C). Importantly, when we limited our analyses to those hybrids with poor piRNA targeting scores (sites with pirScan <-15), we also found that significantly more 22G-RNAs are made from those piRNA targeting sites with CIMs than from those without (Figure 4D). These observations showed that piRNA targeting sites with CIMs elicit more downstream silencing signals. We then evaluated how much WAGO-1 22G-

RNA production at the piRNA targeting sites are contributed by piRNA targeting. If a target site is regulated by piRNAs, we expect a loss of 22G-RNA levels in the *prg-1* mutant, which loses all piRNAs. In CLASH- identified piRNA targeting sites from hybrids not carrying CIMs, we observed a reduction in WAGO-1 22G-RNA levels at piRNA targeting sites. Notably, we found a significantly greater reduction of WAGO-1 22G-RNA levels for those targeting sites from hybrids with CIMs than those without (Figure S3A). Similar observations were found for those hybrids with poor targeting scores (Figure S3B). These observations suggest that CLASH- identified piRNA targeting sites from hybrids with CIMs are in general under stronger control by the piRNA pathway than those without CIMs.

We then compared CLASH-identified miRNA target sites from hybrids with CIMs to those without CIMs for their ability to regulate mRNA expression. Since miRNAs can trigger the mRNA degradation of their targets, we use mRNA levels as a proxy for evaluating the effects of miRNA-mediated gene regulation^29^. In CLASH identified sites without CIMs, we found that target mRNA expression is globally elevated in the *alg-1* mutant. Notably, we found that those miRNA target sites identified through hybrids with CIMs, either deletions or substitutions, exhibit a significantly greater increase in their mRNA levels in the *alg-1* mutant than those without CIMs (Figure 4E). This trend is also found for miRNA sites with poor miRNA targeting scores (miRanda score <140) (Figure 4F).

Together, our analyses showed that RNA target sites identified from CLASH experiments exhibit overall more functional relevance if they are detected from hybrids carrying CIMs. CIM analysis therefore provides a critical tool to identify candidates of functional piRNA and miRNA targeting sites, including those that exhibit non-canonical base pairing. The comprehensive data of hybrid reads and the location of CIMs of C. elegans piRNA and miRNA can be found in supplemental data file 1 (for piRNA) and supplemental data file 2 (for miRNA).

## Discussion

In this manuscript, we developed the MUTACLASH pipeline to identify and analyze CIMs in CLASH data. We examined the location, distribution, and functional implication of CIMs in the miRNA and piRNA CLASH data in *C. elegans*. Our data revealed that CIMs are present in these libraries and that the CIM pattern on mRNAs represents miRNA Argonaute and PIWI footprints on their target mRNAs. Specifically, we found that CIMs are generally enriched at the center region of piRNA and miRNA targeting sites. In addition, CIMs in piRNA CLASH data are also enriched at regions where mRNA and piRNA pairing contain local mismatches. As crosslinking requires zero distance between RNA and certain amino acids, the distribution of CIMs/crosslinking sites provide insight into the interactions between PIWI and Argonaute with their target mRNAs. For example, the enrichment of CIMs at the middle region of PIWI and Argonaute targeting may reflect the relatively narrow channel at the center region for target and mRNA pairing or the proximity between catalytic residues of PIWI/Argonaute and target mRNA nucleotide corresponding to the 10^th^ and 11^th^ nucleotide of small RNAs that can be cleaved upon extensive basepairing^30^. In addition, one model to explain the preference of CIMs in regions with local mismatches is that mismatches between piRNA and mRNA can lead to bulges, and such bulges protruding from mRNA strands, closing the distance between the RNA and PIWI protein and theirfore the chance of crosslinking between PIWI and these nucleotides. Furthermore, as canonical (seed perfect) and non-canonical (seed imperfect) of worm piRNA binding is associated with distinct CIM distribution patterns (Figure 3D), our results suggest that PIWI PRG-1adopts distinct conformations between these two target recognition modes, resulting in distinct CIMs patterns (Figure 5).

**Figure 5.**
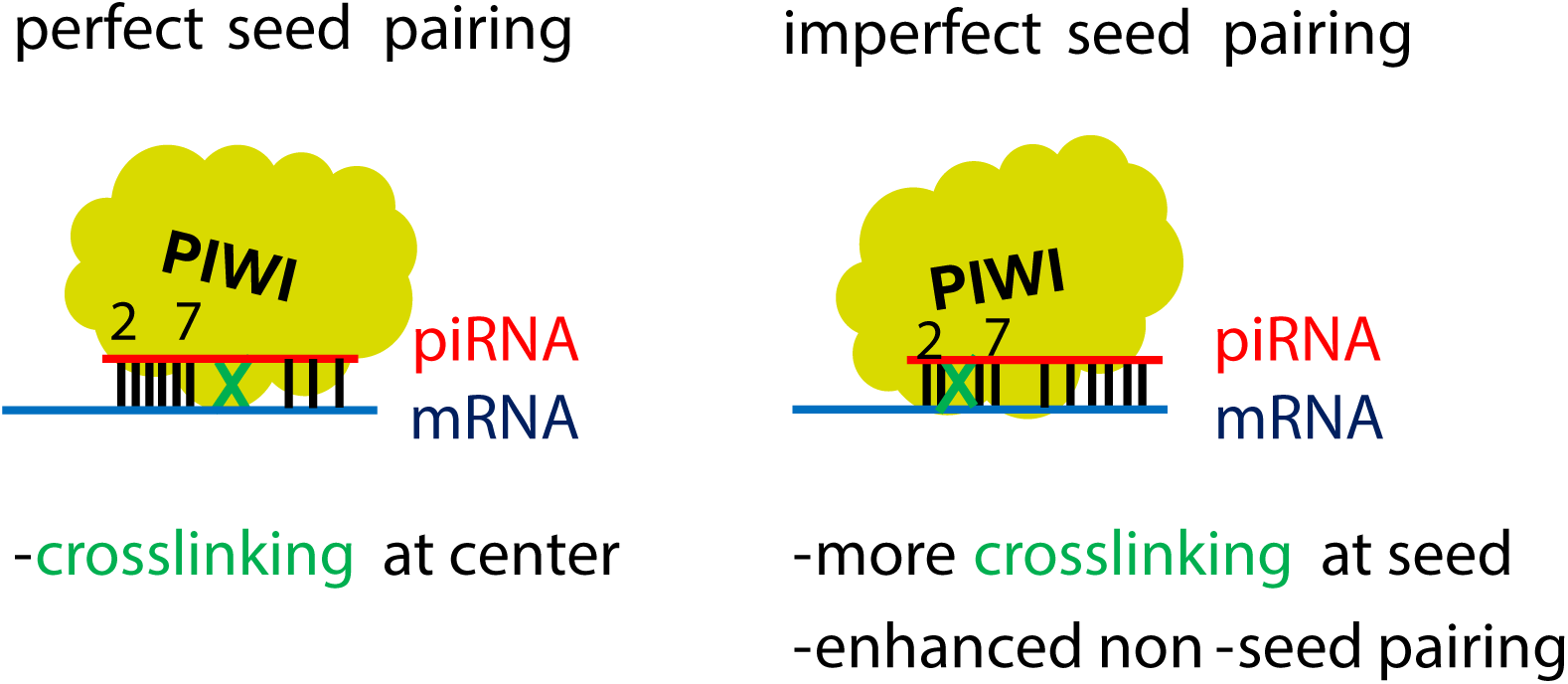
A model depicting the distinct conformations of the PIWI PRG-1 Argonaute protein during target mRNA binding with perfect or imperfect seed pairing, resulting in the enrichment of CIMs at distinct positions.

While reported piRNA targeting rules in *C. elegans* do not tolerate seed mismatches^4^, our analyses of piRNA targeting sites with CIMs suggest that some non-canonical base pairing interactions with seed mismatches are capable of triggering gene silencing. Indeed, few recent studies have suggested that piRNAs can tolerate a seed mismatch in target recognition to trigger gene silencing in *C. elegans* and in other animals^6,23,31^. Notably, despite piRNA targeting sites with CIMs in their seed region exhibiting a decreased pairing ratio at their seed regions, we observed an increased pairing ratio at non-seed regions. This implies that non-seed pairing may complement imperfect seed base-pairing in piRNA target recognitions (Figure 5).

As only a small portion of CLASH data contain canonical small RNA targeting sites, identification of functional small RNA targeting sites from CLASH data have been a major challenge. Our analyses demonstrated that CLASH hybrids with the presence of CIMs exhibit significantly more regulatory effects than those without, including those with poor targeting scores. In addition, while non-hybrid mRNA reads from CLASH data are typically also ignored from previous CLASH data analyses, our CIMs analyses suggest that these non-hybrid mRNAs with CIMs are Argonaute-crosslinked mRNAs *in vivo* (Table S1A). Therefore, both CIMs analyses from hybrid and non-hybrid mRNAs in CLASH data provide valuable information about mRNA targets whose expressions is regulated by small RNAs *in vivo*.

## Methods

### MUTACLASH pipeline

MUTACLASH pipeline can be used to process raw reads from CLASH experiments to identify *in vivo* small RNA binding sites. In brief, the adapter and barcode sequences from the raw reads were trimmed using trim_galore (version 0.6.5) with the following command: --length 17 --dont_gzip -q 30 --max_length 70. Reads were deduplicated, and their read counts were calculated using the custom Python scripts. The hybrid reads consisting of piRNA and mRNA transcripts were identified by ChiRA algorithm^16^ using the following commands: -b -p 4 -l1 12 - go1 6 -mm1 4 -s1 18. The *C. elegans* mRNA transcripts (WormBase WS275 version and piRNA sequences from wormbase (WS275 version and type 2 piRNA sequence^32^ were used as reference sequences. Once the hybrids are identified, the mutated positions on mRNAs were obtained through reading the MD tag with a custom Python script and Samtools^33^. Mutations identified in overlapping regions of mRNA and small RNA sequences are removed to avoid misinterpretation, as these mutations can be derived from either small RNA or mRNAs. The precise location of the mutations on mRNAs are reported. A custom Python script is then used to calculate the nucleotide composition and percentage of each mutation type of CIMs.

### Map the location of CIMs within the predicted small RNA targeting sites

When the mRNA interacting sequences (CLASH identified regions) are shorter than miRNA or piRNA, they are first extended to the size of miRNA and piRNAs using both the upstream and downstream sequences before they are examined for sites with best pairing energy/score. The predicted miRNA or piRNA binding sites are defined with Miranda^34^ or pirScan^21^, respectively. The following commends are used for distinct tools; pirScan commands: --ex n<u>^[d]^</u>, miRanda commands: -sc 0. Custom Python scripts are used for plotting and analysis, including converting the absolute location of mutation into relative location in the small RNA targeting sites, plotting the read count distribution, visualizing the mutation ratio at each position, and generating graphs for the pairing ratio and abundance results.

### Measurements of WAGO-1 22G-RNAs levels at piRNA targeting sites

The *C. elegans* transcriptome data (WS275) annotation was used for mapping *C. elegans* reads. WAGO targets (n=3644) are defined as transcripts whose mapped 22G-RNAs exhibit over two-fold enrichment from either WAGO-1 IP than that from input 22G-RNAs^35^.

For measurements of 22G-RNA reads around piRNA targeting sites, the 50 nt (+/- 25 nt) window centered at the 10^th^ nucleotide of piRNA sequence was used to calculate the read count number of the 22G-RNA reads mapped to these regions.

### mRNA level analysis from RNA seq

Fastq reads were trimmed of adaptors using cutadapt^36^. Trimmed reads were aligned to the C.elegans genome build WS275 using bowtie2 ver 2.3.0^37^. After alignment, reads were overlapped with genomic features (protein-coding genes, pseudogenes, transposons) using bedtools intersect^38^. Reads per kilobase million (RPKM) values were then calculated for each individual feature by summing the total reads mapping to that feature, multiplied by 1e6 and divided by the product of the kilobase length of the feature and the total number of reads mapping to protein-coding genes.

### Datasets

The iCLIP data of ALG-1 (SRR3882949) and the CLASH data of PRG-1 (SRR6512652) were used in these analyses^8,9^. WAGO-1 associated small RNA data (SRR8482951/WT, SRR8482949/*prg-1* mutant) were used for analysis of WAGO-1 22G-RNA levels around piRNA targeting sites^39^. mRNA seq data of wild type (SRX2826535, SRX2826536, SRX2826587) or from the alg-1(gk214) mutant (SRX2826541, SRX2626542, and SRX2826543) were used^29^.

## Acknowledgements

This work is supported in part by the National Science and Technology Council of Taiwan (NSTC 111-2221-E-006-151-MY3 and NSTC 113-2221-E-006-135-MY3) grants to W.-S.W., and the NIH grant R01-GM132457 to H.-C.L.

## Data availability statement

MUTACLASH pipeline and all the custom scripts used in this manuscript can be found at GitHub deposit link: https://github.com/lu1215/MutaCLASH. All sequencing data analyzed in the manuscript are available at NCBI GEO or ENA database. The SRR numbers of sequencing data used in specific analyses are provided in the Method and dataset section.

---

**Figure S1.**
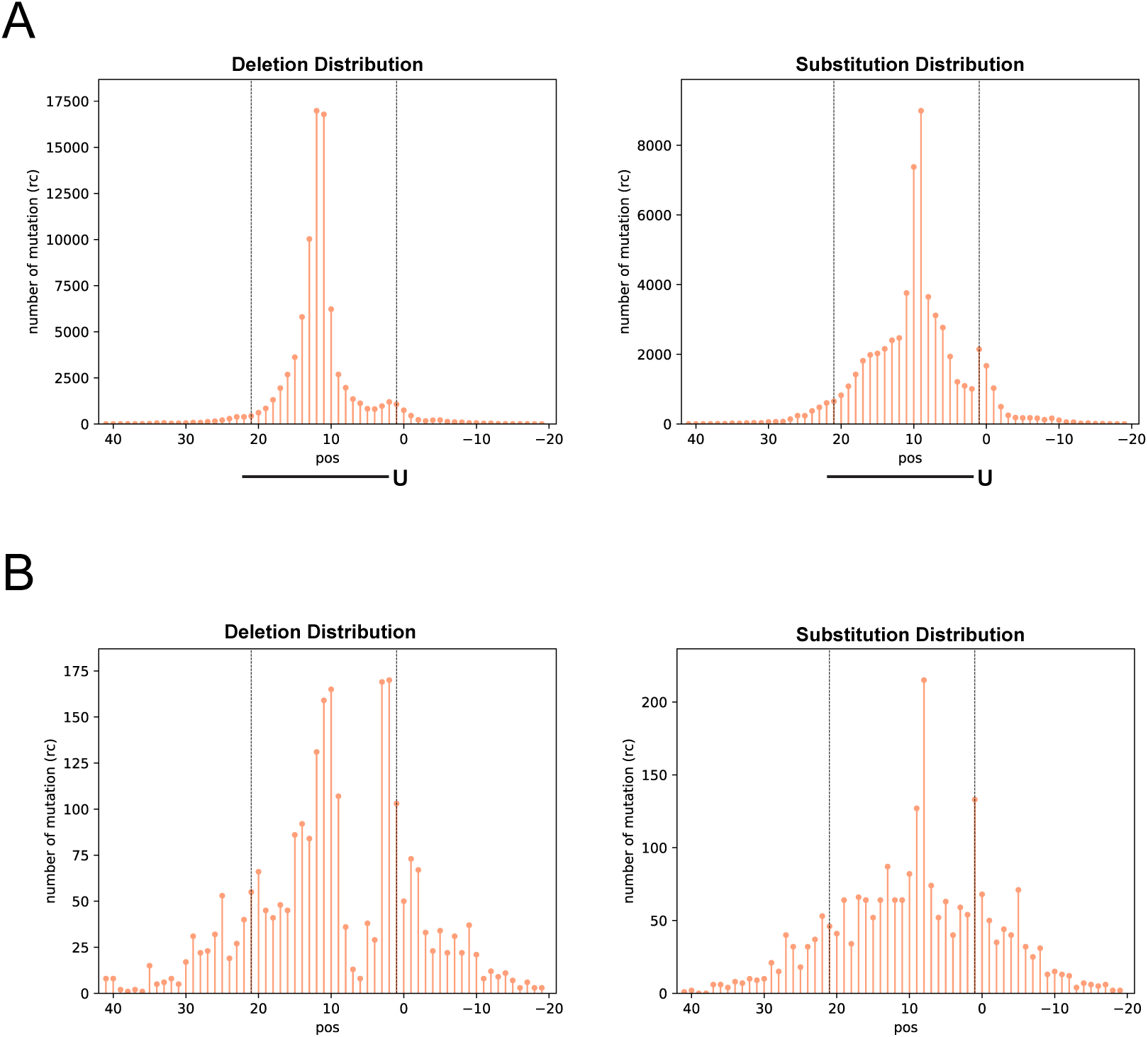
The distribution of CIMs in CLASH Data. A. The number of deletions (left) and substitutions (right) of mRNAs found at the indicated position corresponding to piRNA from those hybrids with the top 33% piRNA targeting score of PRG-1 piRNA PIWI CLASH data. B. The number of deletions (left) and substitutions (right) of mRNAs found at the indicated position corresponding to miRNA from hybrids with top 33% targeting score of ALG-1 miRNA Argonaute CLASH data.

**Figure S2.**
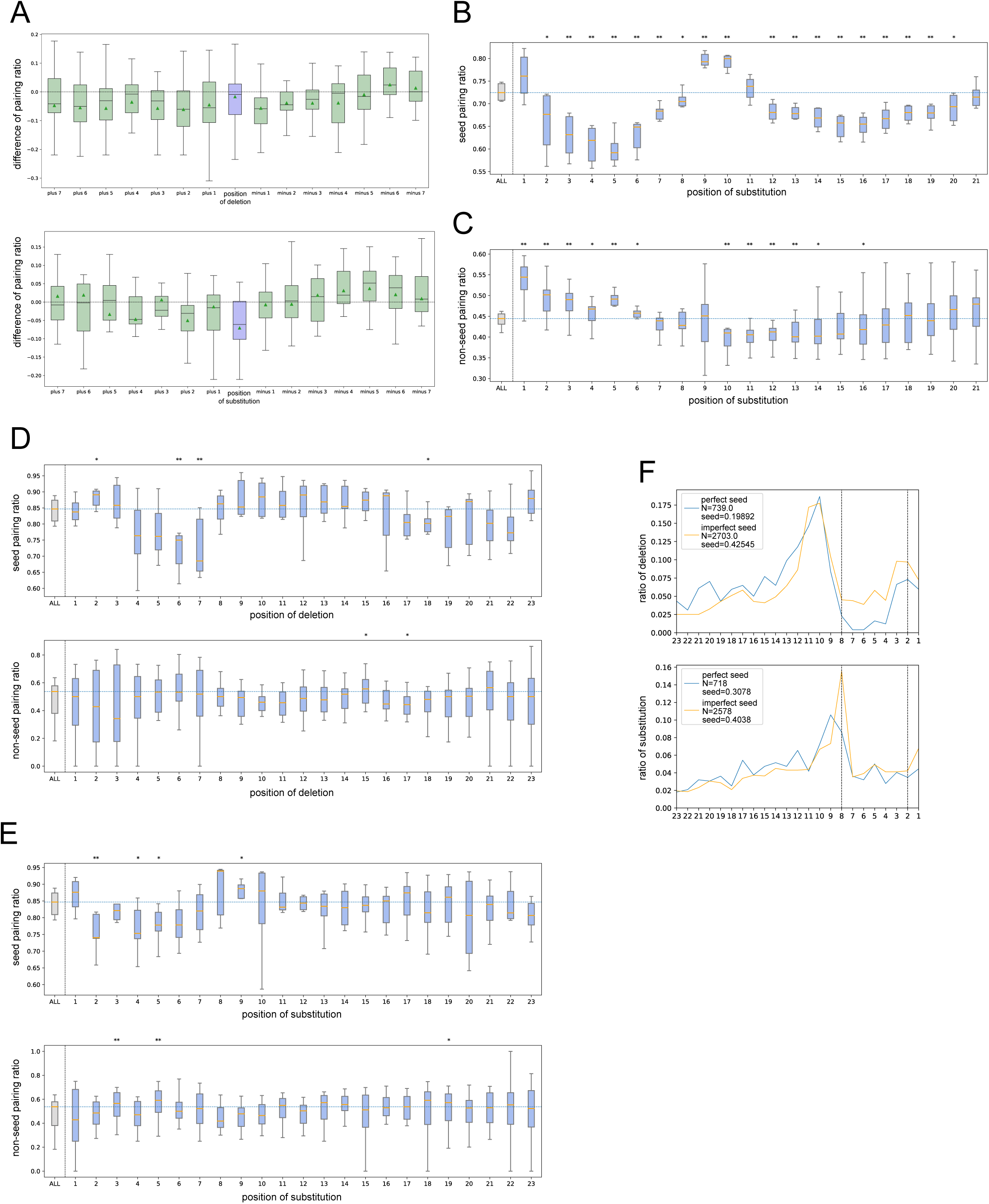
The relationship of CIM with base-pairing ratio in CLASH data. A. Differences in base-pairing ratios around CIMs in ALG-1 miRNA AGO CLASH data. The differences were calculated by subtracting the pairing ratios of hybrids with deletions (top) or substitutions (bottom) from the pairing ratios of all hybrids. The analysis was centered on the CIM location (position 0) and extended to the indicated upstream and downstream nucleotides in ALG-1 miRNA AGO CLASH data. B. Differences of pairing ratio at the seed region between hybrids with substitution at the indicated position and hybrids with CIMs in PRG-1 PIWI CLASH data. C. Differences of pairing ratio at the non-seed region between hybrids with substitution at the indicated position and hybrids with CIMs in PRG-1 PIWI CLASH data. D. Differences of pairing ratio at the seed region (top) or non-seed region (bottom) between hybrids with deletion at the indicated position and hybrids with CIMs in ALG-1 miRNA Argonaute CLASH data. E. Differences of pairing ratio at the seed region (top) or non-seed region (bottom) between hybrids with substitution at the indicated position and hybrids with CIMs in ALG-1 miRNA Argonaute CLASH data. F. Distribution of mRNA deletions (left) and substitutions (right) in hybrids with either perfect seed pairing (blue line) or imperfect seed pairing (orange line) in ALG-1 miRNA Argonaute CLASH data.

**Figure S3.**
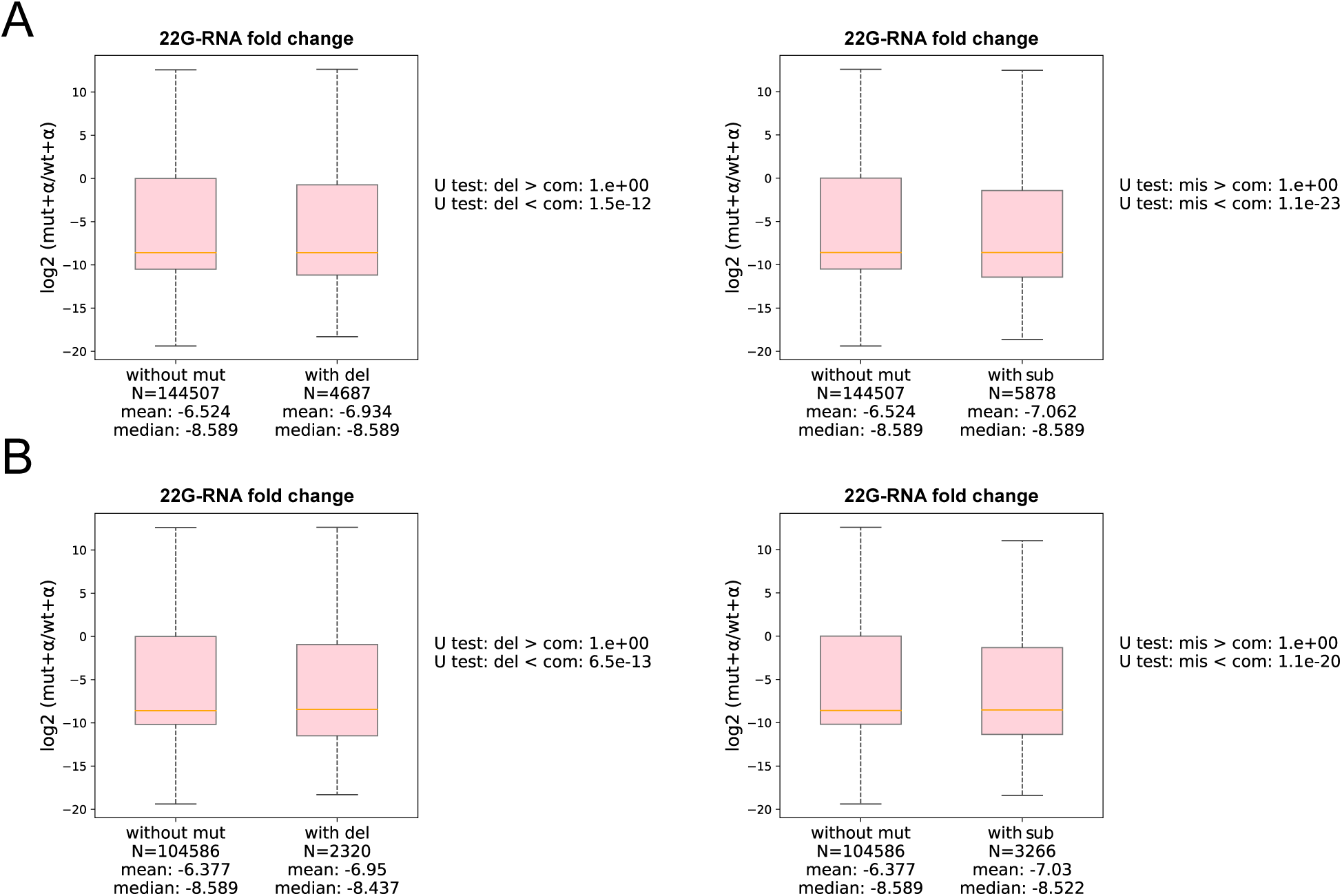
piRNA targets with CIMs exhibit stronger reduction of WAGO-1 22G-RNAs in the *prg-1* mutant. A. The ratio of local WAGO-1 22G-RNAs in the *prg-*1 mutant over those in wild type at piRNA targeting sites without CIMs or at those sites with deletion (left) or with substitution (right). The statistical significance (P-value) of the difference in fold change was calculated by the Mann-Whitney U test. B. The ratio of local WAGO-1 22G-RNAs in the *prg-*1 mutant over those in wild type at piRNA target sites with non-canonical binding sitees (pirScan score < -15) without CIMs or at those sites with deletion (left) or with substitution (right). The statistical significance (P-value) of the difference in fold change (*prg-1* mutant/WT) between two regions was calculated by the Mann-Whitney U test.

**Table S1A.** Frequency of reads with mRNA deletion or substitution in RNA sequencing data or CLASH data.

| Library | Read type | # reads with deletion | # reads with substitution | # total reads | % reads with mutation |
| --- | --- | --- | --- | --- | --- |
| mRNA | Non-chimeric | 5166.5 | 281215 | 20038571 | **Deletion: 0.026%**<br>**Substitution: 1.40%** |
| PRG-1 CLASH | Non-chimeric | 265382 | 270426.5 | 4053251 | **Deletion: 6.55%**<br>**Substitution: 6.67%** |
| ALG-1 iCLIP | Non-chimeric | 875559.5 | 1120123 | 15107177 | **Deletion: 5.8%**<br>**Substitution: 7.41%** |
| PRG-1 CLASH | Chimeric, mRNA | 108970 | 129279 | 4017410 | **Deletion: 2.71%**<br>**Substitution: 3.22%** |
| ALG-1 iCLIP | Chimeric, mRNA | 4897 | 6155 | 133974 | **Deletion: 4.59%**<br>**Substitution: 3.67%** |

**Table S1B.**
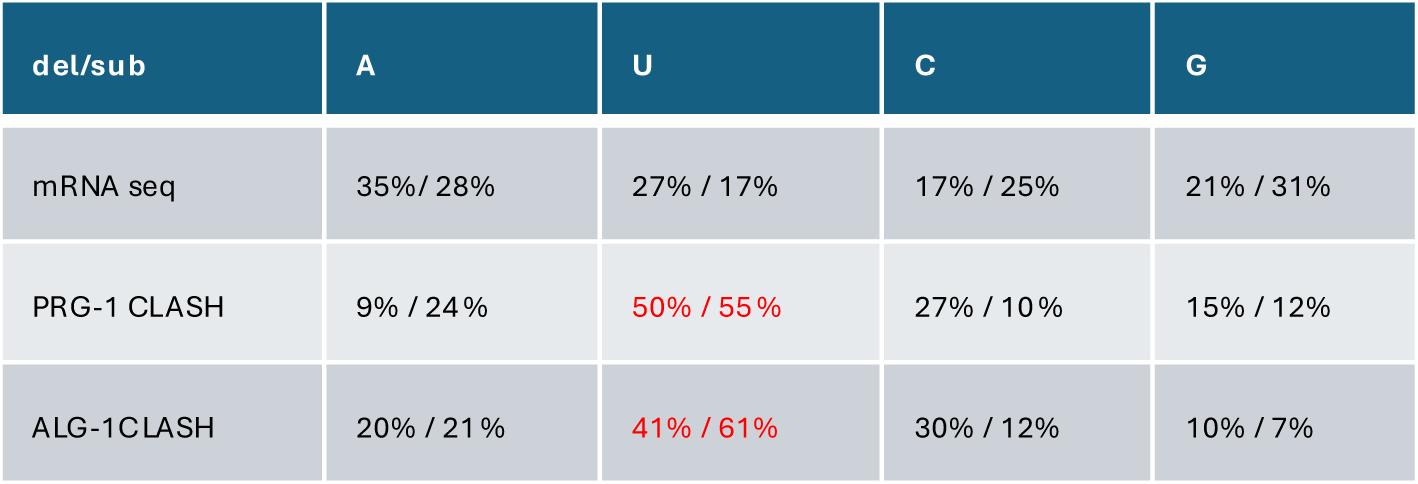
Percentage of the indicated nucleotides that exhibit substitution (sub) or deletion (del) in the indicated dataset.

**Table S1C.**
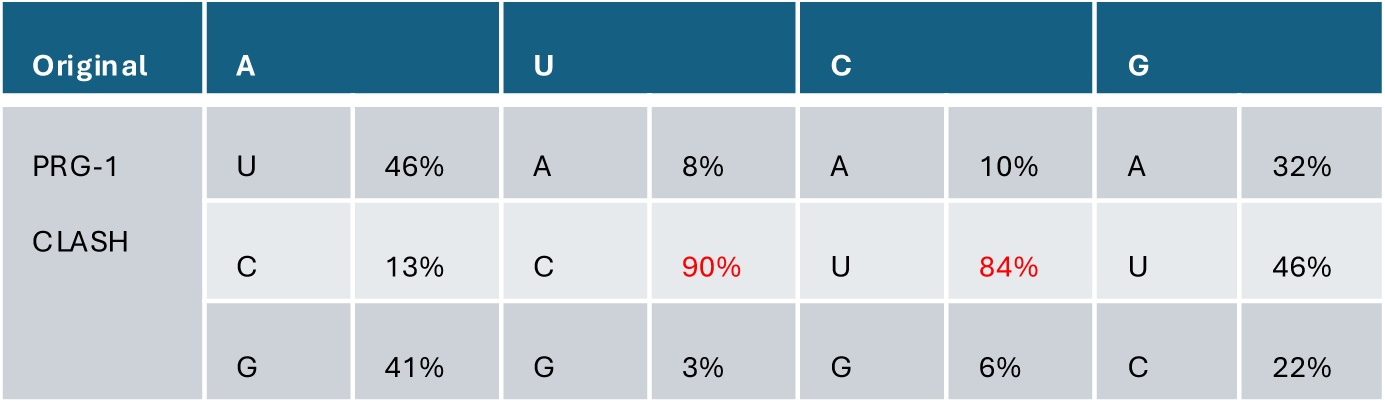
Percentage of the indicated type of mRNA substitutions in the PRG-1 (piRNA) CLASH data.

**Table S1D.**
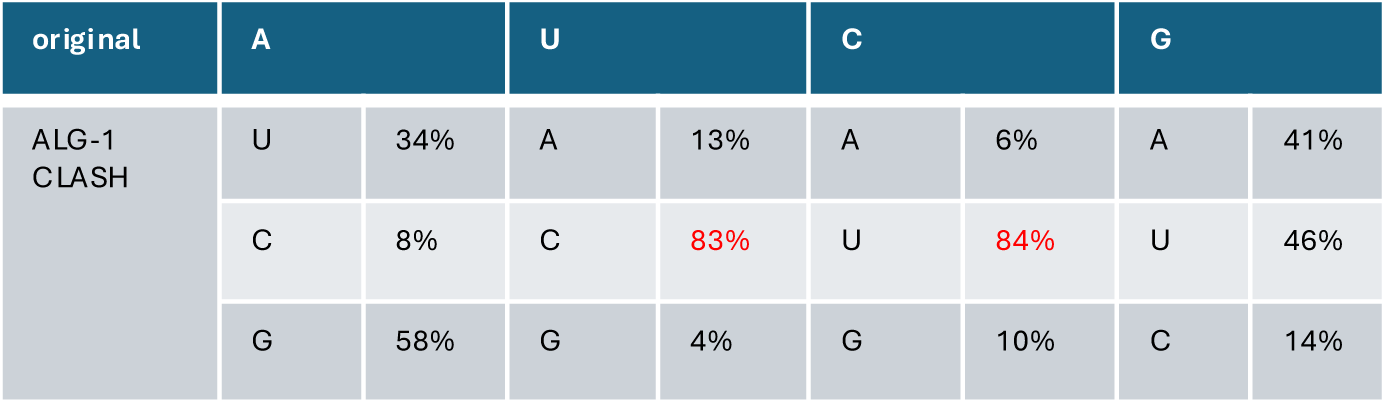
Percentage of the indicated type of mRNA substitutions in the ALG-1 (miRNA) CLASH data.

**Table S2A.** Distance between the location of mRNA deletion and substitutions in PRG-1 piRNA (top) and ALG-1 miRNA (bottom) target CLASH data.

|  |  |  |  |  |  |  |  |  |  |
| --- | --- | --- | --- | --- | --- | --- | --- | --- | --- |
| Position<br>difference | -4 | -3 | -2 | -1 | 0 | 1 | 2 | 3 | 4 |
| # of count | 1443 | 2929 | 5892 | 5421 | 2947 | 1192 | 1001 | 878 | 761 |
| position<br>difference | -4 | -3 | -2 | -1 | 0 | 1 | 2 | 3 | 4 |
| # of count | 670 | 856 | 3265 | 3196 | 1508 | 580 | 427 | 394 | 381 |

